# Assessment of pharmacokinetics, safety, and neuroprotective efficacy of an adjunct intramuscular verapamil therapy in a rat model of organophosphate DFP-induced status epilepticus

**DOI:** 10.1101/2025.06.25.661332

**Authors:** Yam Nath Paudel, Robert E. Blair, Elisa Hawkins, Matthew S. Halquist, Melissa Morgan, Jason Funderburk, Daniel Calvano, Jennifer Koblinski, Hope Richard, Laxmikant S. Deshpande

**Author notes:** To whom correspondence should be addressed: Laxmikant S. Deshpande, M.Pharm., Ph.D., FAES. Virginia Commonwealth University, School of Medicine PO Box 980599, Richmond, VA 23298.

## Abstract

Lethal organophosphate (OP) exposure leads to status epilepticus (SE), which, despite standard-of-care therapy, is associated with acute mortality and long-term morbidities. Neuronal injury and inflammation are reported following OP-SE, and drugs targeted at these processes have produced beneficial outcomes. Verapamil (VPM) is a calcium-channel blocker used as an antihypertensive drug. It exhibits neuroprotective and anti-inflammatory actions in experimental models of CNS injuries. Here, we investigated the feasibility of an adjunctive intramuscular (i.m.) VPM therapy in OP Diisopropyl Fluorophosphate (DFP)-induced SE. We also investigated the safety and toxicity of i.m. VPM and compared its pharmacokinetic (PK) profile to oral (p.o.) administration. Rats were injected with DFP (4 mg/kg, s.c.). One minute later, atropine (0.5 mg/kg, i.m.) and 2-PAM (25 mg/kg, i.m.) were injected, and at 1-hour post-SE, midazolam (1.78 mg/kg, i.m.) was administered. Rats were then treated with VPM (10 mg/kg, i.m, 3 days). Histological analysis was conducted to assess neuronal injury and injection-site pathology. In a separate group of rats, PK studies were conducted on blood and brain homogenates treated once with saline or VPM (10 mg/kg, p.o. or i.m.). Our data demonstrated that following DFP-SE, i.m. VPM achieved higher blood and brain levels and exhibited a favorable PK profile compared to p.o. route. VPM therapy did not cause significant muscle pathology and produced a robust neuroprotective response. Our studies provide evidence that the i.m. route is an effective method for delivering VPM following SE producing significant neuroprotective outcomes compared to treatment with the standard-of-care alone in OP-SE.

## Introduction

Status Epilepticus (SE) is a clinical emergency associated with high mortality and significant neurological morbidities^1–3^. There are several causes for SE, including stroke, traumatic brain injury, withdrawal from anti-seizure medications, alcohol withdrawal, and exposure to certain toxic chemicals ^4^. One such category of chemicals that could lead to the rapid induction of SE is the organophosphate (OP) compounds^5, 6^. These compounds are diverse in their applicability and include commonly used pesticides, industrial chemicals, and additives to jet fuels^7^. OP compounds also include lethal chemical warfare nerve agents (CWNA)^8^. OP exposure can thus occur accidentally, occupationally, domestically, or during war/ terrorism-related scenarios^9–12^. Furthermore, intentional use of OP compounds for self-harm is also unfortunately common in developing nations ^13, 14^.

OP compounds are inhibitors of the enzyme acetylcholinesterase, leading to accumulation of acetylcholine at the synapse^8, 15, 16^. Depending on the OP dose and the level of AChE inhibition, symptoms of OP exposure range from salivation, lacrimation, and headaches to tremors, muscle fasciculations, and seizures. These symptoms could further evolve into unremitting seizures (status epilepticus, SE), respiratory difficulties, and eventually death if prompt medical care is not provided^17, 18^. Emergency treatment for OP intoxication includes respiratory support and treatment with a three-drug regimen standard of care (SOC) recommended by the Food and Drug Administration (FDA) to manage hypercholinergic signs. These drugs include atropine (a muscarinic receptor antagonist), pralidoxime (an AChE reactivator), and midazolam (a benzodiazepine to manage seizures)^19, 20^. These emergency procedures have significantly reduced the acute mortality associated with OP exposures. However, despite these treatments, survival from OP exposure is associated with chronic neurological co-morbidities, including memory impairments, mood disorders, and even spontaneous recurrent seizures (SRS). Thus, effective treatments are needed to control not only the acute mortality but also lower the incidences of chronic morbidity associated with OP exposures ^21, 22^.

Cellular injury and neurodegeneration are key processes underlying the neurological consequences of OP toxicity ^15, 23–25^. Our lab has studied neuronal calcium homeostatic mechanisms in rat models of OP toxicity^26^. We reported protracted elevations in hippocampal calcium levels using the paraoxon (POX)^6^ and diisopropyl fluorophosphate (DFP)^5^ models of OP-induced SE. Calcium-induced calcium release mechanisms were found to mediate these chronic elevations in calcium. Indeed, treatment with ryanodine receptor (RyR) and inositol-tris-phosphate (IP3R) antagonists such as dantrolene and levetiracetam not only lowered the OP-SE-induced neuronal calcium levels but also produced a neuroprotective effect^27^. Ketamine, which prevents calcium entry by blocking the N-methyl-D-aspartate receptors (NMDAR), has also been neuroprotective in some OP-SE models^28, 29^. Neuroinflammation has also been reported in OP-SE, and pharmacological treatment aimed at suppressing neuroinflammatory processes is being investigated to improve neurological outcomes following OP intoxication^30–35^.

Verapamil (VPM) is a phenylalkylamine calcium channel blocker approved by the Food and Drug Administration (FDA) in the 1980s for the management of high blood pressure as well as for treating angina and cardiac arrhythmias ^36^. The non-FDA-approved indications for VPM include its use for treating cluster headaches ^37^, acute coronary syndrome ^38^, and cerebral vasospasm ^39^. The drug is typically administered orally, and an intravenous formulation is available. The usual dose range is 120 to 360 mg/day given orally and up to 10 mg parenterally^40^. VPM has a long human safety record with minimal side effects and drug-drug interactions ^41^. Recent findings regarding the molecular actions of VPM, in addition to calcium-channel inhibition, have demonstrated a potent neuroprotective and anti-inflammatory action, raising the possibility that VPM could also be an effective OP countermeasure drug. For example, VPM increased neuronal survival and memory in a mouse stroke model ^42^. VPM (20 mg/kg, i.p.) also prevented neuronal damage and rescued memory following severe hypoglycemia in rats ^43^. VPM (10 mg/kg, i.p.) is also reported to decrease apoptosis in a rat model of ischemia/ reperfusion injury ^44^. In a cellular model of Alzheimer’s Disease, destabilization of calcium homeostasis and enhanced glutamate excitotoxicity were noted, which were attenuated by VPM ^45^. Similarly, VPM was reported to be neuroprotective in a cellular model of dopaminergic neurotoxicity ^46^. VPM also rescued motor neurons by reducing endoplasmic reticulum stress in a mouse model of ALS ^47^. Together, these findings point toward a neuroprotective profile for VPM, providing a rationale for testing VPM as a potential therapy to attenuate OP-SE toxicities.

In this study, using a rat model of DFP-induced SE^5^, we compared the pharmacokinetics of VPM when administered orally (p.o.) or intramuscularly (i.m.). The i.m. route was selected given the necessity for ease of administration during an OP-based mass casualty scenario. VPM is also a substrate for P-glycoprotein (P-gp), a transmembrane protein that moves drugs out of the cell ^48, 49^. Therefore, we also measured brain levels under both SE and no-SE conditions to investigate the brain penetrance and availability of VPM. We also investigated the neuroprotective efficacy of adjunctive i.m. VPM therapy in various brain regions following DFP-induced SE when co-administered with the standard FDA-approved three-drug regimen of atropine, 2-PAM, and midazolam indicated for OP intoxication. Muscle pathology at the injection site was also evaluated to investigate the safety of repeatedly administering i.m. injections. VPM therapy. Finally, a stability analysis of our VPM formulation was also conducted.

## Materials and Methods

### Drugs and Chemicals

DFP (purity > 90%) was obtained from Chem Service Inc. (West Chester, PA). Verapamil hydrochloride, atropine sulfate, and pralidoxime chloride (2-PAM) were obtained from Millipore-Sigma (St. Louis, MO). Midazolam vials (5 mg/mL) were obtained from Med-Vet International (Mettawa, IL). To prepare a working DFP solution (4 mg/mL), the desired quantity of DFP was removed using a Hamilton syringe and added to a glass bottle containing ice-cold phosphate-buffered saline (PBS), and then gently vortexed. The bottle was kept on ice, and syringes were drawn and kept on ice until the time of subcutaneous (s.c.) injections. Time on ice (between dilution in PBS and injection) was never more than ten minutes. Verapamil, Atropine sulfate, and 2-PAM were dissolved in saline (SAL, 0.9% NaCl) and then sterile-filtered.

### Animals

All animal use procedures were per the National Institute of Health Guide for the Care and Use of Laboratory Animals and approved by Virginia Commonwealth University’s Institutional Animal Care and Use Committee. Male and Female Sprague-Dawley rats were obtained from Envigo (Indianapolis, IN) at 9 weeks of age. Animals were housed two per cage at 20-22°C with a 12 Light: 12 Dark hour cycle (lights on 0600-01800 h) and free access to food and water.

### Induction of DFP-SE

As shown in Figure 1, in accordance with our previously published protocol for inducing DFP-induced SE^5, 50^, separate cohorts of rats were injected with DFP (4 mg/kg, s.c.). One minute later, atropine (0.5 mg/kg, i.m.) and 2-PAM (25 mg/kg, i.m.) were injected to permit survival and progression into SE. DFP exposure produced signs of progressive cholinergic hyperstimulation culminating in continuous seizures. We utilized the Racine criteria for assessing SE severity^51, 52^. The criteria for the behavioral seizure score were: 0 - behavioral arrest, hair raising, excitement, and rapid respiration; 1 - mouth movement of lips/tongue, vibrissae movement, and salivation; 2 – head "bobbing"/clonus; 3 – forelimb clonus; 4 – clonic rearing, and 5 – clonic rearing with loss of postural control. A Racine score of 4-5 was used as the criterion to mark the onset of SE, approximately 5-10 minutes post-DFP injection. Our previous studies have shown that this Racine score correlates with the tonic-clonic phase of SE and is accompanied by an EEG frequency of >6 Hz, which meets the clinical criteria of SE^5, 50^. At 1-h post-SE onset, midazolam (1.78 mg/kg, i.m.) was administered to suppress SE. Midazolam administration was delayed by 1 h post-DFP-SE onset to simulate a practical therapeutic window for first responders or hospital admission following mass casualty events^5, 53, 54^. These doses were determined using human equivalents calculated using a human-to-rat dose translation equation^55, 56^. Control rats were age-matched, no-SE condition that received either SAL or VPM.

**Figure 1:**
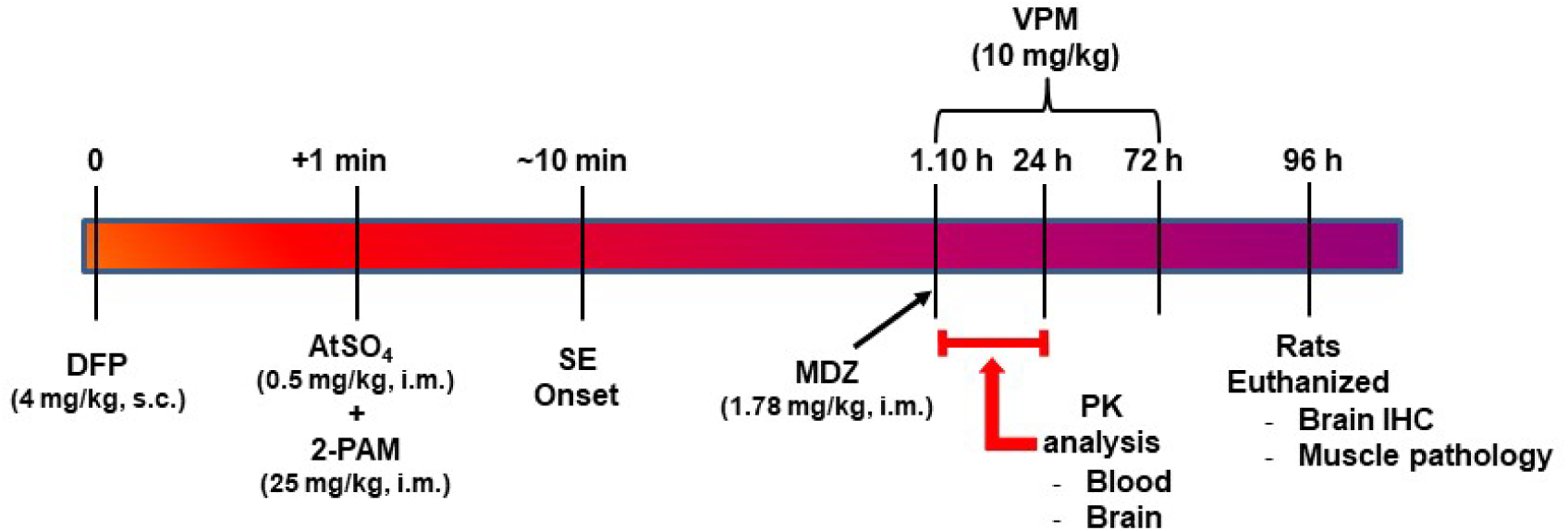
Timeline for induction of DFP-SE and VPM interventions post SE. At time 0, Sprague-Dawley rats (both male and female) were injected with the OP agent DFP (4 mg/kg, s.c.). One minute later, atropine sulfate (AtSO_4_; 0.5 mg/kg, i.m.) and 2-PAM (25 mg/kg, i.m.) were injected. SE onset was approximately 10 min. Rats were allowed to seize for 1 h, at which point midazolam (1.78 mg/kg, i.m.) was injected. For PK studies, VPM (10 mg/kg) was administered once via the p.o. or i.m. route, and blood and brain samples were collected at various time points up to 24 hours. For histological studies, rats were injected with saline or VPM (10 mg/kg, i.m., *b.i.d.*) for three consecutive days. Mortality was assessed daily till the fourth day, when the rats were euthanized and perfused for brain and muscle collection.

### Verapamil treatment

Rats that met the SE severity criteria (Racine 4-5) and received midazolam at 1-h post-DFP-SE onset were randomized into two treatment groups: those receiving saline (SAL) or VPM. Previous studies indicated that VPM could be administered i.p. in rodent seizure models at doses up to 40 mg/kg ^57–59^. A pilot study with two doses of VPM (10 and 20 mg/kg, i.m.) at 1 h after DFP-SE was conducted in rats (n=5). Rats safely tolerated both doses in these pilot studies. For this study, we chose a 10 mg/kg twice-daily treatment regimen as this dose was closer to the human-equivalent dose ^56^ and would allow for a stable plasma level given a t_1/2_ of 5 h in humans ^60, 61^. Thus, VPM was administered concomitantly with the first dose of midazolam at 60 minutes post-DFP-SE onset and subsequently twice daily for up to 72 hours, for a total of six VPM doses. The control group was given i.m. SAL injections for the same duration and frequency. For i.m. administration, VPM or SAL was administered in the quadriceps muscle using a 28-gauge needle. The injection volume was 0.1mL/kg. Injection sites were alternated between the lateral and contralateral sides for the twice-daily dosing protocol. Manifestations of distress and discomfort (such as limping, licking the injection site, and other ambulation-related issues) were subjectively graded as: none, mild, moderate, and severe. Mild discomfort was noted on the last day of i.m. injections, which quickly resolved without any intervention. No ambulatory deficits were observed in these rats. Muscle pathology was conducted as described below.

### Pharmacokinetic analyses

For the pharmacokinetics study, rats were dosed once with SAL or VPM (10 mg/kg). Blood samples were obtained in duplicate from 3-4 rats at each of the 10 time points for the two routes of administration (i.m. and p.o.) following saline or DFP administration. Following anesthesia, rats were euthanized and blood samples were collected through cardiac ventricles at 0, 5 min, 30 min, and at 1, 2, 4, 6, 12, 18, and 24 h following VPM administration. The blood was placed in an EDTA microcuvette and frozen at -80 °C until analyzed. Concurrently with blood collection, the brain was rapidly dissected, and the cortex was removed and snap frozen in isopentane (-15 ^°^C) and stored at -80 ^°^C until use. VPM levels were measured in whole blood and cortex using liquid chromatography tandem mass spectrometry (LC-MS/MS). The linear range was 0.05-1200 ng/mL for VPM.

### Sample preparation

50 µL of rat whole blood was analyzed for VPM levels using a protein precipitation extraction method. Briefly, each aliquot was transferred to a 1.5 mL micro-centrifuge tube, and 500 µL of extraction solvent consisting of 70% methanol and 30% water was added, following the 50 µL of internal standard solution (AT-d 7 and LV-d 6 at a concentration of 500 ng/mL). Sample tubes were vortexed for 1 min and sonicated for 30 min. Samples were then centrifuged for 10 min at 13.2g and transferred to an amber vial. VPM was analyzed using LC-MS/MS with multiple reaction monitoring (MRM) transitions on an AB Sciex 6500+ Qtrap (Sciex, USA) with an LC-20-AD (Shimadzu, USA) UPLC system. Chromatographic separation was performed on a Zorbax Eclipse Plus C18 Column (2.1 X 100mm, I.D. 3.5 µm, Agilent Technologies, USA). The column temperature was ambient; the flow rate was 1.0 mL/min, and the injection volume was 1 µL. The mobile phase A consisted of 0.2% formic acid in water, and mobile phase B in methanol. Isocratic conditions were used at acetonitrile (95%) and 0.2% formic acid (5% ) in reagent water. Data were analyzed with Analyst 1.6 Quantitation Wizard in accordance with the Bioanalytical Laboratories Standard Operating Procedures.

### Perfusion and tissue collection

At 96-h post DFP-SE, a separate group of rats underwent perfusion and fixation procedures to obtain brain and muscle tissue. Following induction of deep anesthesia with ketamine/xylazine (75 mg/ 7.5 mg/kg, i.p.), rats underwent transcardial perfusion with isotonic saline and then perfused with 250 ml of 4% paraformaldehyde in a 0.1 M phosphate buffer (pH 7.4). Brains and quadricep muscles were removed and allowed further fixation for 24 h in buffered 4% paraformaldehyde at 4 ^°^C. Brains were then cryoprotected in 30% sucrose in 0.1 M phosphate buffer (pH 7.4) for 3 days at 4 ^°^C. Brains were then embedded in OCT by snap freezing in isopentane (-15° C) and stored at -80°C until use. Post-fixed muscle was transferred to a 70% ethanol solution and processed as described below.

### Fluoro-Jade staining

Coronal brain sections were prepared using a Leica CM3050S cryostat (Leica Biosystems, Deer Park, IL) and adhered to glass slides (Superfrost Plus; Fisher Scientific, Pittsburgh, PA) and stored at -80 ^°^C until use. Briefly, 20 μm-thick sections were collected between (-1.8 mm to -4.56 mm Bregma, based on rat anatomic atlas ^62, 63^). The sectioning spans the entire dorsal hippocampal formation and includes the somatosensory cortex, thalamus, amygdala, and piriform cortex. Each slide contained sections representing all experimental groups (age-matched control, DFP-SAL, DFP-VPM). Every 20th section through the rat hippocampus is selected from at least six animals for labeling and staining ^27, 53, 54, 64–66^.

Slides were dried in a desiccant chamber at 55 °C for 30 min prior to staining. Slides were first incubated in a solution of 1% NaOH in 80% ethanol for 5 min, followed by hydration in 70% ethanol and then ddH_2_O for 2 min each. Slides were then incubated in a 0.06% KMnO_4_ solution for 10 min, followed by washing in ddH_2_O for 2 min. Slides were then stained in a 0.0004% Fluoro-Jade C (FJC) solution in 0.1% acetic acid for 20 minutes. Stained slides underwent 3× washes in ddH_2_O for 2 min each and then dried in a desiccant chamber at 55 °C for 30 min. Stained slides were then cleared with xylene for 5 min and cover-slipped with DPX mounting agent. FJC-stained sections were viewed under 20X magnification on an Olympus IX-70 microscope under brightfield or fluorescein/FITC filter settings, and 16-bit grayscale digital images were acquired with a digital CCD camera (ORCA-ER; Hamamatsu Corp) using MetaMorph7.8 (Molecular Devices, San Jose, CA). All microscope/camera settings for FJC image acquisition were constant throughout. Grayscale images were analyzed using ImageJ 1.53 (NIH). Specific staining was measured using auto-threshold segmentation (Yen method) of selected regions of interest and represented as cell counts/mm^2^ (analyze particles module)^27, 65, 66^.

### Assessment of muscle pathology

To evaluate the safety of i.m. VPM administrations, quadricep muscle specimens were obtained at 1 day and 3 months post the last i.m. injection of the VPM treatment regimen. To conserve animal numbers, muscle specimens for acute toxicity assessment were obtained from the same rats that were utilized for dissecting the brain for FJC staining. A separate cohort of rats was utilized for the 3-month time point. These time points were selected to assess whether repeated VPM injections caused any muscle injury at the site of injection, either acutely or chronically. Muscle specimens were obtained as described in the perfusion section above. Muscle samples were embedded in paraffin, and 5-micron-thick sections were obtained. These sections were then stained with Hematoxylin and Eosin (H&E) for histopathological analysis. Automated H&E staining was performed with the Dako CoverStainer (Agilent). A standard four-point inflammation scale was used for scoring ^67, 68^ as described in Table 1. The score for each site was the average of the scores for each of the three tissue blocks.

**Table 1:** Inflammation scale for assessment of injection site muscle pathology.

| **Score** | **Fibrosis** | **Infiltrates** |
| --- | --- | --- |
| 0 = Not present | 0 | None present |
| 1 = Minimal | <1 mm | Few, widely scattered infiltrates |
| 2 = Mild | 1 to 1.9 mm | Scattered infiltrates, minimal small clusters |
| 3 = Moderate | 2 to 3.9 mm | loosely packed fields, many small clusters |
| 4 = Marked | >4 mm | Dense infiltrates, often packed fields |

### Stability analysis of VPM formulation

VPM formulation (10 mg/mL in sterile saline) in multiple glass bottles was stored at 25°C ± 2°C and 60 ± 5% Relative Humidity. At the respective time points (Day 0, 90, 203), three bottles were randomly chosen, and the stability of the VPM formulation was assessed under ICH QA1 (R2) guidelines [Stability Testing of New Drug Substances and Products] and using a similar USP VPM injection monograph (USP29-NF24).

Similar to the USP29-NF24 VPM monograph, a system suitability solution was prepared at 1.9 mg/mL of USP Verapamil Hydrochloride RS and 1.5 mg/mL of USP Verapamil Related Compound B RS in mobile phase (see Table 2). The VPM standard solution (2.5 mg/mL) was also prepared in the mobile phase. Sample Preparation (Verapamil HCL Injection Sample): Dilute sample to a working solution of 2.5 mg/L in mobile phase. Dilute working solution to 0.5 mg/L in the mobile phase. Impurity Standard solution: 2.5 mg/mL of USP Verapamil Hydrochloride RS, and 7.5 µg/mL each of USP Verapamil Related Compound A RS, USP Verapamil Related Compound E RS, and USP Verapamil Related Compound F RS in Mobile phase. Filter the samples and standards using a syringe filter into 1 mL autosampler tubes.

**Table 2:** HPLC Conditions for VPM concentration determination.

|  |  |
| --- | --- |
| Column | 4.6-mm ×15-cm; 5-μm packing L1 |
| Mobile Phase | Acetonitrile, 0.015 N Sodium Acetate, 3.3% acetic acid, and 2-aminoheptane (30:70:0.5) |
| Needle Wash | 60:30:10 Acetonitrile:Methanol:Water |
| Flow Rate | 0.9 mL/min |
| Injection Volume | 10 μL |
| Detector | UV 278 nm, Range 200-400 nm for Identification |
| Column Temperature | 30°C |
| Sample Temperature | 6°C |
| Run Time | 4 min (NLT 4 times retention time of verapamil) |
| Software | Waters Empower |

### Experimental rigor and Data analysis

Male and Female rats were housed in separate rooms. All the drugs were prepared fresh daily and belonged to the same respective batch to minimize experimental variability due to possible minor differences between drug lots. DFP-SE induction protocol and VPM treatment were carried out on separate days for male and female rats. Rats were randomized, ignoring sex and the estrous cycle stages. Experimental groups were blinded until the data were analyzed. The study excluded animals that did not present with SE onset or failed to achieve a Racine score of 4 or higher. Two independent observers assessed the severity of SE. Data analysis was performed using GraphPad Prism 10 software. Data represented as mean ± SD. Mixed-effect analysis utilized a two-way ANOVA. Multiple t-tests were employed as appropriate. A p-value ≤ 0.05 was considered to indicate statistical significance.

For pharmacokinetic analysis, model-independent methods were used to estimate the pharmacokinetic parameters of VPM in rat blood samples. The area under the curve (AUC) was determined utilizing the parameters programmed in Microsoft Excel. AUC from time zero to the last sampling time was calculated using the linear trapezoidal rule. The AUC from the last sampling time was extrapolated to infinity by dividing the last measured plasma concentration by the terminal elimination rate constant. Cmax and Tmax were obtained directly from the concentration-time curves. The AUC data are expressed as mean ± SD and compared using a One-Way Analysis of Variance (ANOVA) followed by a post-hoc Tukey test with a significance set at p<.05.

## Results

### Pharmacokinetic studies after i.m. and p.o. treatment with VPM

VPM blood levels were measured in control rats (no DFP-SE) at various time intervals after administration by i.m. and p.o. routes. Table 3 presents data on the pharmacokinetics (PK) of VPM treatment in control rats. In both control male and female rats, increased bioavailability (AUC) and higher drug concentrations in plasma (Cmax) were observed when VPM was administered i.m. compared to p.o. The time to peak VPM level (Tmax) was delayed following i.m. dosing under control conditions.

**Table 3.** Summary of i.m. to p.o. pharmacokinetics for Verapamil in control rats.

| **Verapamil (males)** |  |  |  |  |  |
| --- | --- | --- | --- | --- | --- |
|  | **AUC<sub>trap</sub>**<br>**[ng/ml\*min]** | **AUC<sub>∞</sub> [ng/ml\*min]** | **t<sub>max</sub>**<br>**[min]** | **t<sub>1/2</sub>**<br>**[min]** | **C<sub>max</sub>**<br>**[ng/ml]** |
| IM | 153774 | 18717055 | 60 | 159 | 994 |
| PO | 16591 | 2109810 | 32 | 389 | 379 |
| **Verapamil (females)** |  |  |  |  |  |
|  | **AUC<sub>trap</sub>**<br>**[ng/ml\*min]** | **AUC<sub>∞</sub> [ng/ml\*min]** | **t<sub>max</sub>**<br>**[min]** | **t<sub>1/2</sub>**<br>**[min]** | **C<sub>max</sub>**<br>**[ng/ml]** |
| IM | 117482 | 18316706 | 50 | 152 | 773 |
| PO | 27597 | 6376058 | 22 | 248 | 213 |

Verapamil (10 mg/kg) was administered to control rats separately via intramuscular (i.m.) or oral (p.o.) route. Animals were sacrificed at various time points after injection, and blood was collected for analysis. Descriptive pharmacokinetic parameters are C<sub>max</sub>, maximal plasma concentration, where plasma concentration is in ng/ml; T<sub>max</sub>, time to reach C<sub>max</sub>; t<sub>1/2</sub>, terminal half-life; AUC<sub>trap</sub>, area under the plasma concentration time curve 0→∞ via trapezoidal method; AUC<sub>∞</sub>, area under the plasma concentration time curve extrapolated to infinity.

Next, we investigated the PK profile of i.m. VPM administered after DFP-SE. As shown in Table 4, non-compartmental analysis revealed comparable PK values for various PK parameters in male and female rats. However, compared to the control (no-DFP SE) condition, an even higher Cmax was obtained after DFP-SE. Furthermore, the time to Tmax after i.m. VPM administration in DFP-SE was decreased compared to control, no-SE rats, and was similar to the values obtained in non-SE conditions upon p.o. administration.

**Table 4.** Summary of i.m. pharmacokinetics for verapamil in Control and DFP-SE rats.

| **Verapamil (DFP, males)** |  |  |  |  |  |
| --- | --- | --- | --- | --- | --- |
|  | **AUC<sub>trap</sub>**<br>**[ng/ml\*min]** | **AUC<sub>∞</sub>**<br>**[ng/ml\*min]** | **t<sub>max</sub>**<br>**[min]** | **t<sub>1/2</sub>**<br>**[min]** | **C<sub>max</sub>**<br>**[ng/ml]** |
| IM- control | 153774 | 18717055 | 60 | 159 | 994 |
| IM- SE | 223027 | 33138419 | 32 | 136 | 1763 |
| **Verapamil (DFP, females)** |  |  |  |  |  |
|  | **AUC<sub>trap</sub>**<br>**[ng/ml\*min]** | **AUC<sub>∞</sub>**<br>**[ng/ml\*min]** | **t<sub>max</sub>**<br>**[min]** | **t<sub>1/2</sub>**<br>**[min]** | **C<sub>max</sub>**<br>**[ng/ml]** |
| IM- control | 117482 | 18316706 | 50 | 152 | 773 |
| IM- SE | 285769 | 110757434 | 50 | 139 | 708 |

Verapamil (10 mg/kg) was administered to DFP-SE rats via the intramuscular (i.m.) route.
Animals were sacrificed at various time points after injection, and blood was collected for analysis. Descriptive pharmacokinetic parameters are C<sub>max</sub>, maximal plasma concentration, where plasma concentration is in ng/ml; T<sub>max</sub>, time to reach C<sub>max</sub>; t<sub>1/2</sub>, terminal half-life;
AUC<sub>trap</sub>, area under the plasma concentration time curve 0→∞ via trapezoidal method; AUC<sub>∞</sub>, area under the plasma concentration time curve extrapolated to infinity.

Figure 2 shows a comparison of plasma concentration (ng/mL) versus time profiles for VPM in male and female rats for the two dosing routes. Mixed-sex analysis indicated that the i.m. route provided significantly higher and sustained blood VPM levels in both male and female rats (Fig. 2A). Significantly higher VPM levels were reached within 30 minutes of i.m. injection and remained significantly elevated at all subsequently measured time points compared to p.o. dosing in both the male (Fig. 2B) and the female (Fig. 2C) rats (n=3-4 rats/sex/ timepoint, multiple t-test for each time point, *p<0.05). No significant differences in VPM levels were noted within the sexes for i.m. or the p.o. routes (Fig. 2D, E).

**Figure 2:**
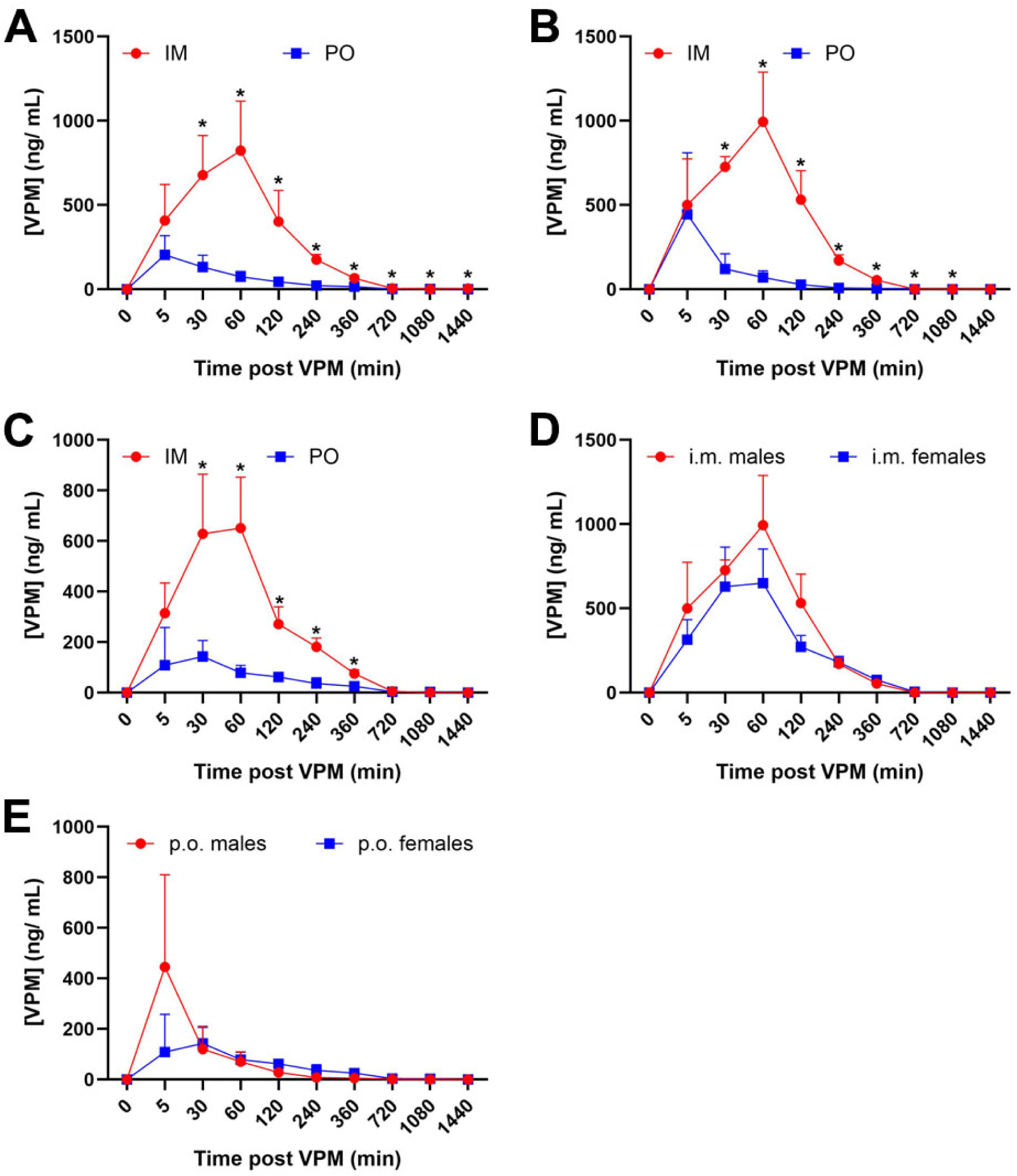
Blood levels of VPM following p.o. and i.m. administration. Comparison of the PK profile of VPM (10 mg/kg) following p.o. and i.m. administration in control rats. Levels of VPM were analyzed using LC-MS/ MS in blood specimens collected from control, no-SE male and female rats at time points ranging from 0 min to 1440 min (24 h). **A.** significantly higher and sustained levels of VPM were noted after i.m. compared to p.o. dosing. **B, C.** significantly higher VPM levels were achieved in both male and female rats. **D, E.** No sex differences were noted for VPM levels following i.m. or p.o. administration. (Each data point represents the mean ± SD of concentration values (ng/ml) from 3 rats. *p<.05, Tukey post-hoc analysis was used to compare concentrations).

### Brain levels of VPM

Upon i.m. administration, VPM was detected in the brains of male and female rats under control and DFP-SE conditions (Table 3). Mixed-sex analysis revealed differences in VPM brain levels under the control and DFP-SE conditions. Following DFP-SE, male rats attained significantly higher VPM concentrations as early as 30 minutes, and this level was sustained for up to 1 hour. Subsequently, male rats continued to exhibit higher VPM concentrations for up to 4 hours. Beyond this time point, the VPM concentration in male rats began to decline while remaining stable in female rats. Significantly higher VPM levels were observed in female rats at the 6-hour time point. At the 18-hour time point, VPM was undetectable in male rats but measurable in female rats (Figure 3; n=3-4 rats/timepoint/group, *p<.05, two-way ANOVA, Bonferroni’s multiple comparison test).

**Figure 3:**
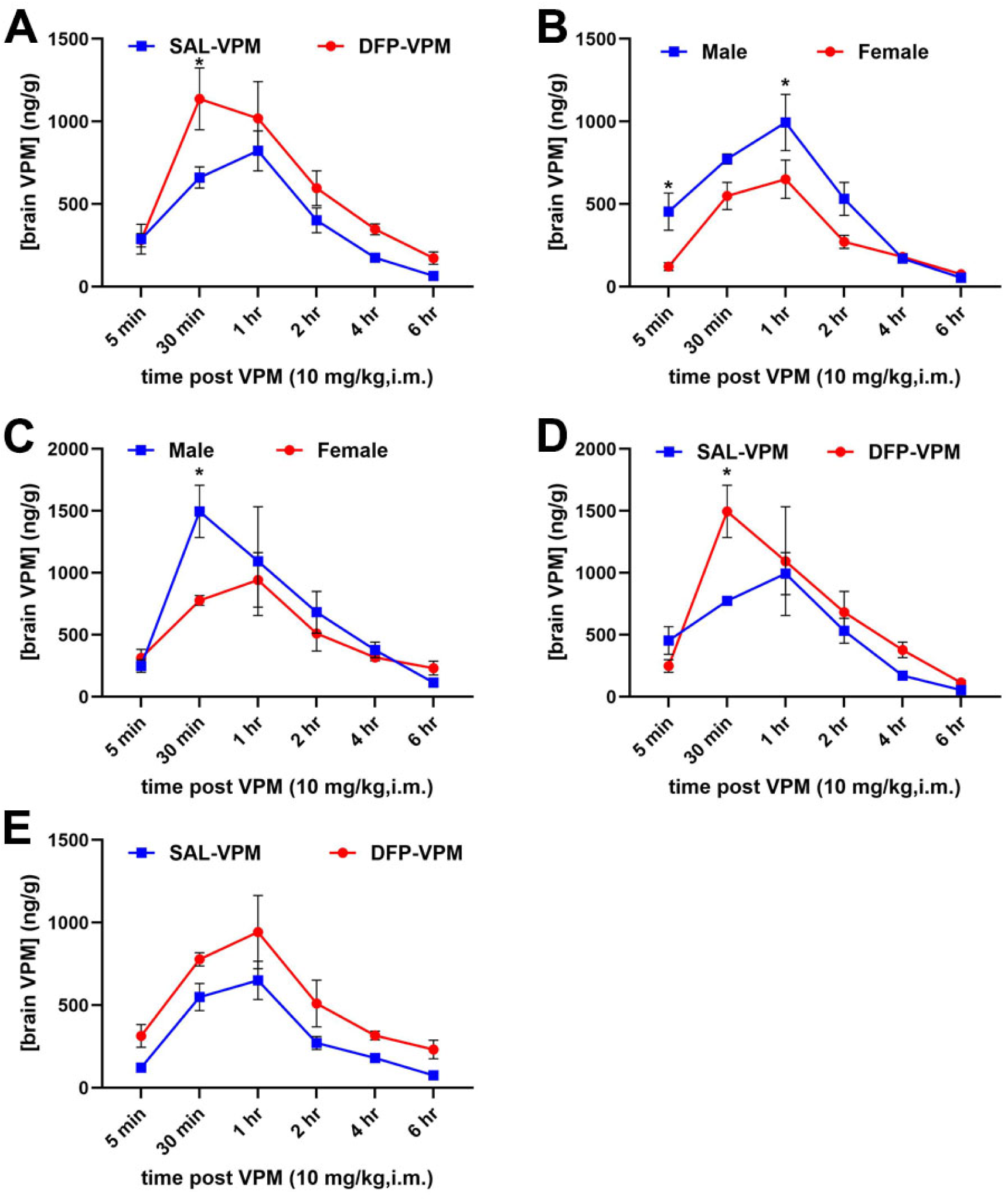
Brain levels of VPM following i.m. administration. VPM (10 mg/kg, i.m.) was administered once to control and DFP-SE male and female rats, and the brain was dissected at various time points ranging from 0 to 24 hours. The concentration of cortical VPM was then estimated using LC-MS/ MS. **A.** Mixed-sex analysis showed higher levels of brain VPM in both control and DFP-SE conditions. **B-C.** Higher levels of VPM were observed under the DFP-SE condition compared to the no-SE condition in both male (B) and female (C) rats. **D-E.** Male rats showed higher VPM levels, which peaked earlier than those in female rats under both control (D) and DFP-SE conditions (E). Significant differences at each time point are indicated (n=3-4 rats/timepoint/group, *p<.05, two-way ANOVA, Bonferroni’s multiple comparison test).

### Muscle safety studies after i.m. treatment with VPM

As illustrated in Figure 4A, one day after the last i.m. injection, the pathological scores for quadricep muscle inflammation were not significantly different between DFP-SE rats treated with SAL and those treated with VPM. As illustrated in Figure 4B, three months post-injections, the muscle pathology score for the DFP-SE rats treated with SAL control did not differ significantly from the VPM tissue. No gender-specific differences were noted in inflammation scores between SAL and VPM treatment at both the acute and chronic time points (n=4 rats/group, two-way ANOVA, *p<.05).

**Figure 4:**
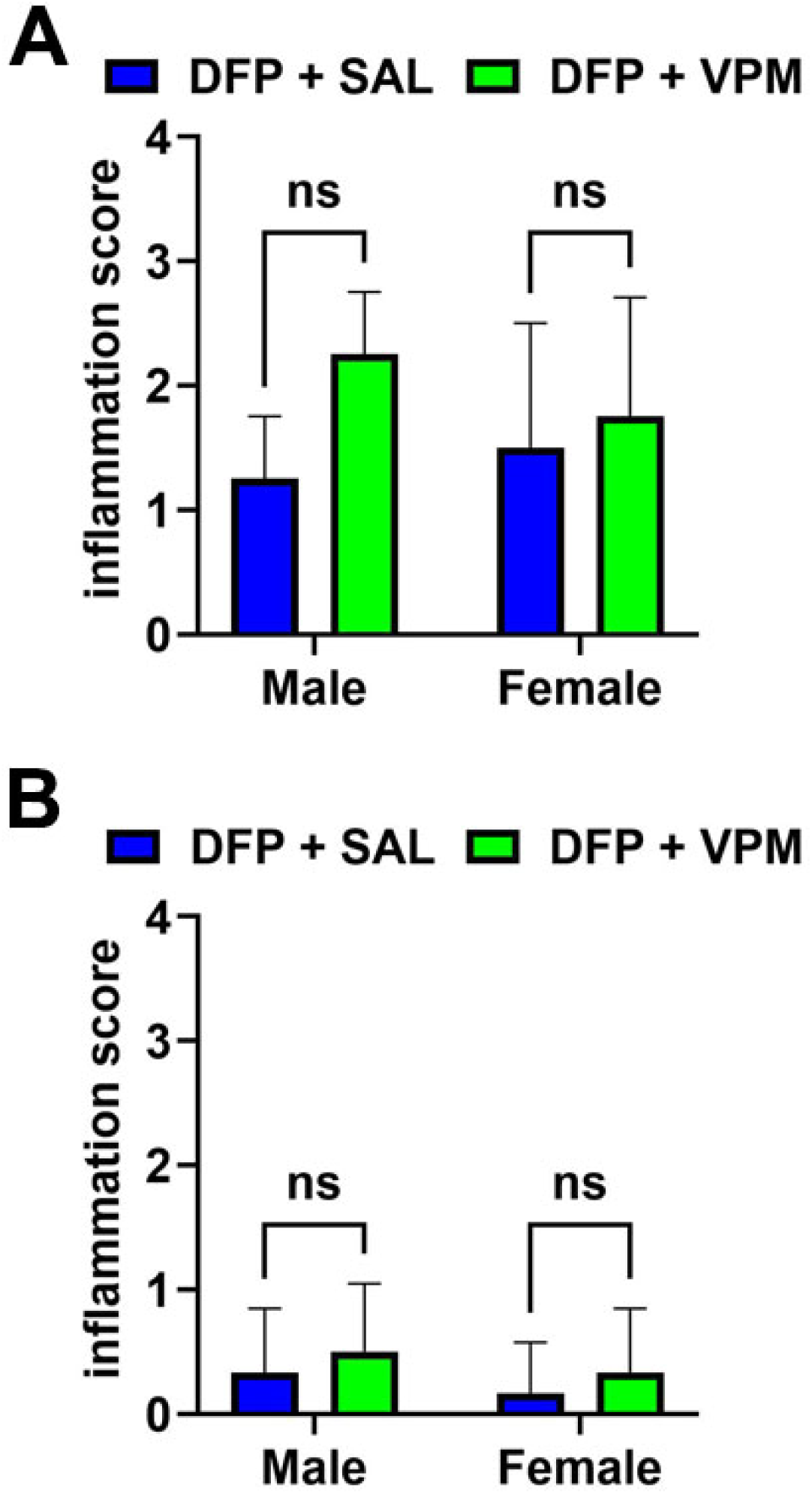
Acute and chronic muscle pathology following i.m. VPM therapy. The data represent the muscle damage score for the i.m. VPM therapy protocol (twice-daily for 3 days) when evaluated 1 day and 3 months after the end of the injections regimen in the following groups: DFP + SAL (blue) and DFP + VPM (green) in male and female rats. No significant sex or treatment differences in the inflammation scores were noted **(A)** acutely or (B) chronically (data represented as mean inflammation score ± SD; n=4 rats/group, two-way ANOVA, *p<.05).

### Assessment of the neuroprotective efficacy of i.m. VPM

Injured brain cells undergoing neurodegeneration in DFP-SE rats exhibited bright green fluorescence in FJC-stained sections (Figure 5A). Sex-specific spatial differences in DFP-induced neuronal injury were observed (Figure 5B-E). In the male rats, significantly more FJC+ cells were found in the CA1, DG, and PC regions compared to female rats (Figure 5E; Two-way ANOVA, post-hoc Tukey test, *p<0.05, n=6-8 rats per group).

**Figure 5:**
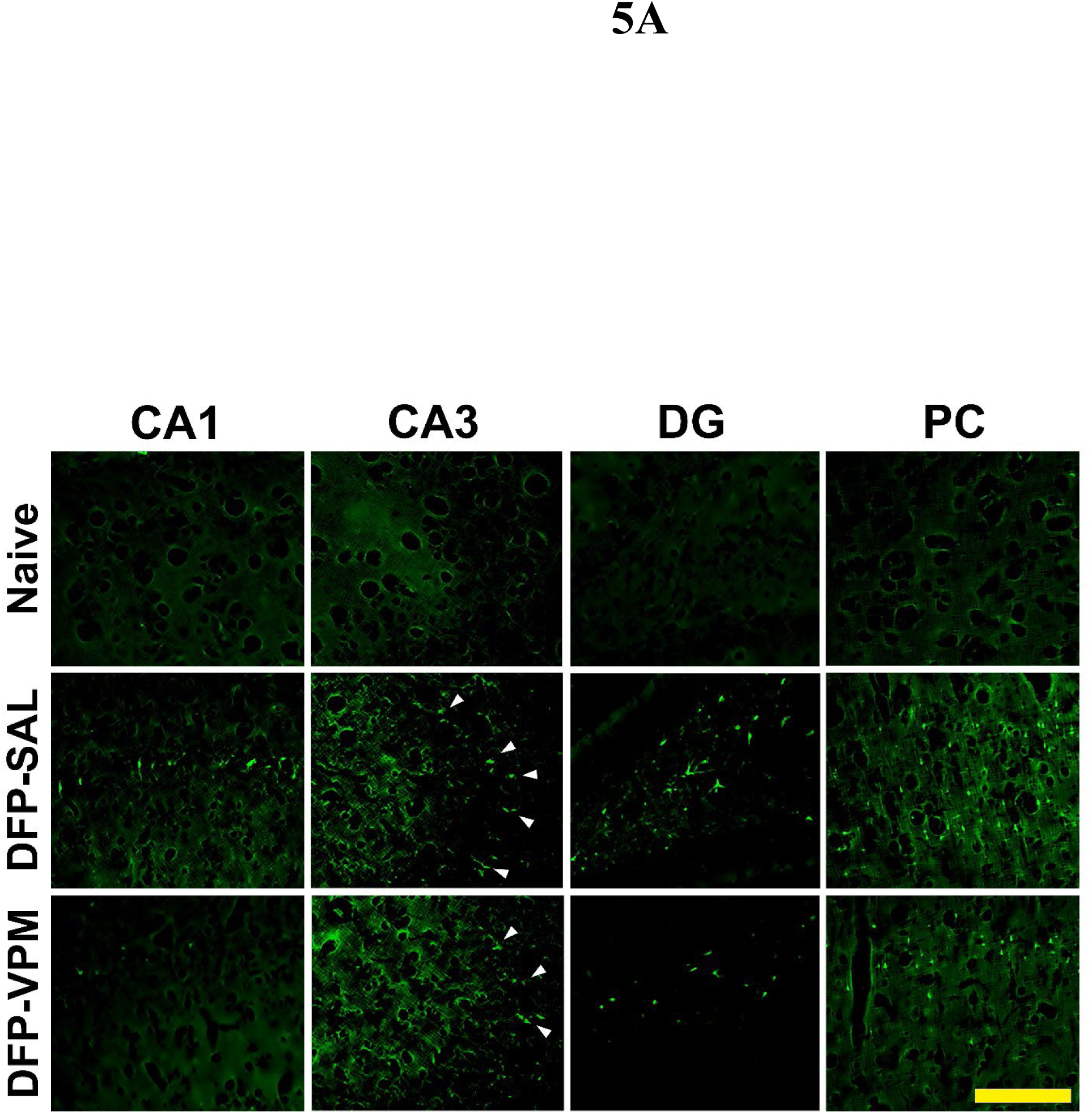
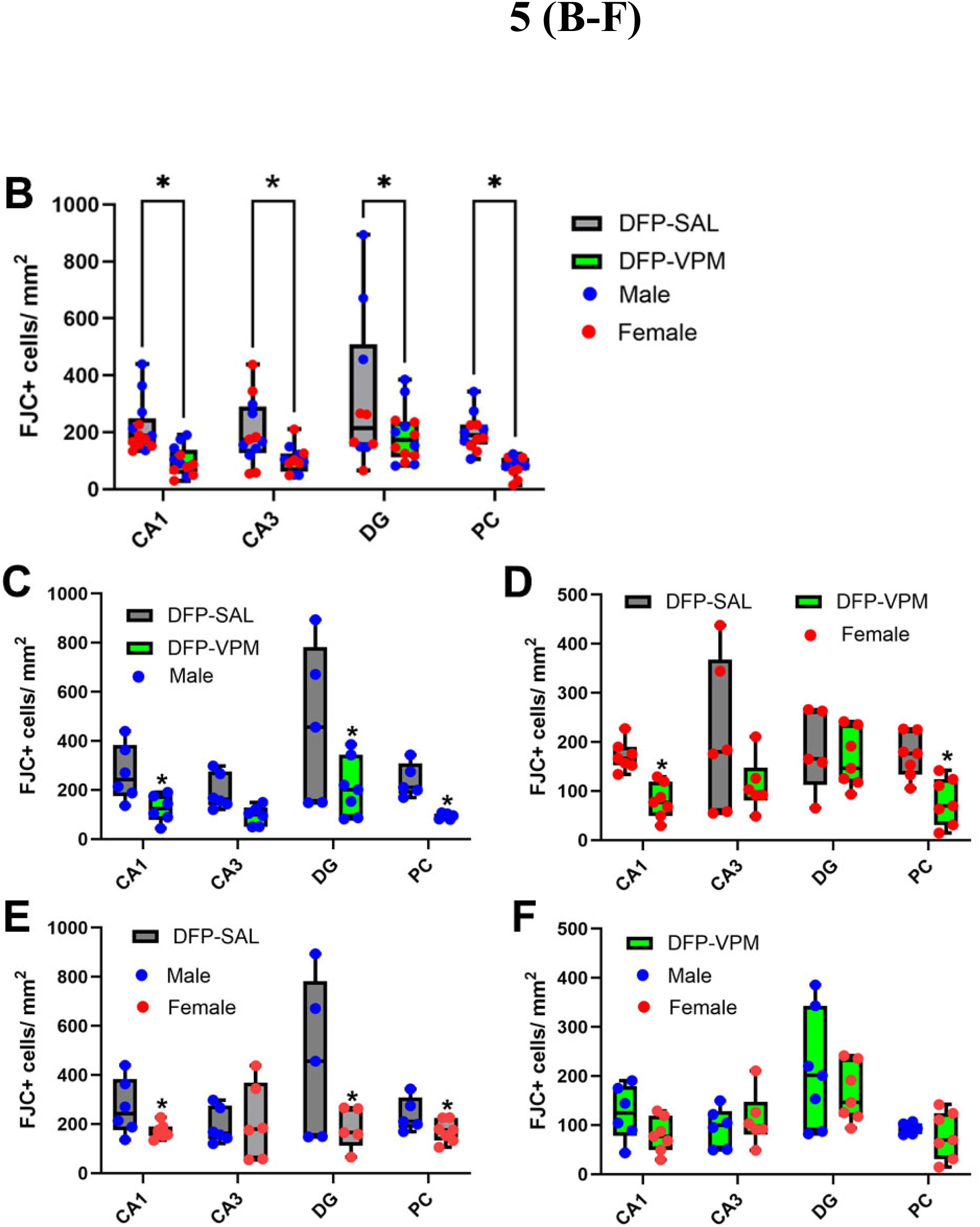
Neuroprotective effects of VPM. **A.** Extensive FJC+ staining indicative of dying neurons is seen in multiple brain regions from DFP-SE rats, which is markedly reduced in VPM-treated DFP rats (scale 200 μM). **B.** A significant decrease in FJC+ cell counts was seen across the brain regions measured in VPM-treated DFP rats compared to SAL-treated DFP rats. **C-D.** Regional differences in VPM’s neuroprotective efficacy were noted between male and female rats following DFP-SE. **E.** Male rats were more sensitive to injury following DFP-SE across the CA1, DG, and PC regions. **F.** No sex differences were noted for the neuroprotective actions of VPM across all the brain regions quantified. (n= 8 per group, *p<.05, two-way ANOVA, Sidak multiple comparison test).

Mixed-sex analysis also revealed that significantly more FJC+ cells were present in the CA1, CA3, and DG regions of the hippocampus, along with the PC, in DFP rats treated with SAL compared to those treated with VPM (Figure 5B-D; Two-way ANOVA, post-hoc Tukey test, *p<0.05, n= 6-8 rats per group). No sex differences in the neuroprotective effects of VPM were observed (Figure 5F). No FJC+ cells were found in brain sections from control rats. (Figure 5A)

### Stability of VPM formulation

Stability studies based on the USP monograph for VPM injection demonstrated that our VPM injectable formulation (10 mg/mL in sterile saline) remained within 90-110% of the labeled claim at all tested time points (0, 90, and 203 days), indicating minimal degradation of VPM in our formulation even after 6 months post-constitution (Table 6, n= 3 bottles assayed at each time point).

## Discussion

The results from our study demonstrate that i.m. VPM can achieve higher blood and brain levels and faster kinetics compared to p.o. administration in both male and female rats. Our VPM formulation had a long shelf life and underwent slight degradation even at 200 days after constitution. Adjunctive treatment regimen consisting of i.m. VPM (10 mg/kg) administered along with the OP-SE standard-of-care therapy, starting 1 hour after DFP-SE onset and continuing for three days, was found to be safe and did not cause significant muscle pathology either acutely or chronically. Finally, i.m. VPM therapy significantly reduced neuronal injury in multiple brain regions following DFP-SE in both male and female rats.

Since this neuroprotective adjunct treatment will initially require on-field administration, an i.m. route of administration is desirable as it provides speed for rapid intervention during a mass exposure event or battlefield-related forward care situations ^69^. VPM is commonly administered p.o. We therefore first compared the PK profile of VPM in whole blood when administered p.o. and i.m. to control, no-SE rats (Table 3). Non-compartmental analyses revealed that VPM, when administered via the i.m. route, produced significantly higher drug levels and exhibited greater bioavailability than when administered p.o. in both male and female rats. In contrast, time to peak VPM level (Tmax) was delayed following i.m. dosing under control conditions. However, following DFP-SE, a higher Cmax was obtained compared to the Cmax following i.m. administration in control, no-SE conditions. Interestingly, the duration to Tmax obtained after i.m. administration in DFP-SE was shorter than the Tmax obtained after i.m. administration in no-SE rats and was comparable to the values obtained in no-SE conditions upon p.o. administration. Overall, these results indicated that the i.m. route provided significantly higher VPM levels, achieved stable therapeutic blood levels, and demonstrated a favorable PK profile, suggesting it could be an effective route to deliver VPM in OP-SE (Table 4; Figure 2).

The blood-brain barrier (BBB) is a diffusion barrier that regulates substances entering the brain. In addition, P-glycoprotein (P-gp) is a transporter protein that protects the brain from xenobiotics. These mechanisms play a role in limiting the entry and levels of drugs in the brain. In a rat model of ischemia-induced CNS injury, the level of VPM penetrance in the brain was reported to be dependent on BBB breakdown and also regulated by P-gp^70^. Thus, we studied VPM brain levels following DFP-induced SE. It has been previously reported that following DFP-SE, the BBB becomes leaky and increased permeability is noticed as early as 6 h following SE and persists for up to 7 days ^71, 72^. Furthermore, VPM is reported to bind and inhibit P-gp directly. In our studies, under both control and DFP-SE conditions, VPM was measurable in the brain upon i.m. administration (Table 5; Figure 3). Higher VPM brain levels were observed in both male and female rats following DFP-SE compared to the control condition. Overall, there was a trend toward greater and quicker VPM brain accumulation in males compared to females. However, a statistical significance was not noted. Interestingly, VPM levels were sustained longer in female rats than male rats under both SE and no-SE conditions. These differences in VPM brain kinetics could partially explain the sex differences seen in the neuroprotective efficacy of VPM. It may also warrant optimization of dose and frequency of administration between the two sexes. Future studies in our lab are planned to investigate these sex differences.

**Table 5.**
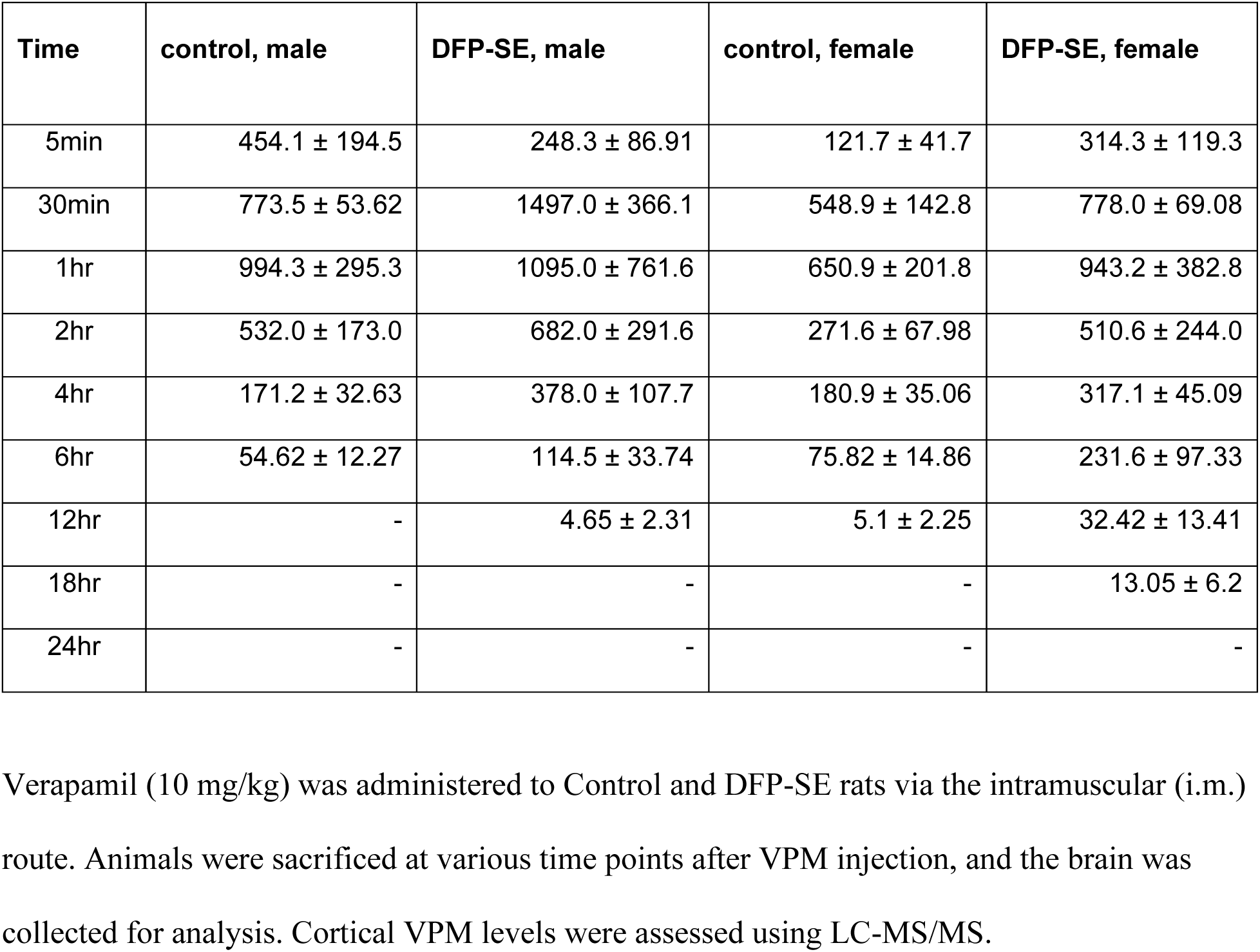
Brain levels of VPM (ng/g) in Control and DFP-SE rats.

| **Time** | **control, male** | **DFP-SE, male** | **control, female** | **DFP-SE, female** |
| --- | --- | --- | --- | --- |
| 5min | 454.1 ± 194.5 | 248.3 ± 86.91 | 121.7 ± 41.7 | 314.3 ± 119.3 |
| 30min | 773.5 ± 53.62 | 1497.0 ± 366.1 | 548.9 ± 142.8 | 778.0 ± 69.08 |
| 1hr | 994.3 ± 295.3 | 1095.0 ± 761.6 | 650.9 ± 201.8 | 943.2 ± 382.8 |
| 2hr | 532.0 ± 173.0 | 682.0 ± 291.6 | 271.6 ± 67.98 | 510.6 ± 244.0 |
| 4hr | 171.2 ± 32.63 | 378.0 ± 107.7 | 180.9 ± 35.06 | 317.1 ± 45.09 |
| 6hr | 54.62 ± 12.27 | 114.5 ± 33.74 | 75.82 ± 14.86 | 231.6 ± 97.33 |
| 12hr | - | 4.65 ± 2.31 | 5.1 ± 2.25 | 32.42 ± 13.41 |
| 18hr | - | - | - | 13.05 ± 6.2 |
| 24hr | - | - | - | - |

Verapamil (10 mg/kg) was administered to Control and DFP-SE rats via the intramuscular (i.m.) route. Animals were sacrificed at various time points after VPM injection, and the brain was collected for analysis. Cortical VPM levels were assessed using LC-MS/MS.

Significant neuronal damage has been reported following OP-SE^5, 6, 30, 33, 64, 73^. Brain regions that show the most severe damage include the hippocampal regions (CA1, CA3, DG), as well as cortical regions such as the parietal cortex and the piriform cortex, along with nuclear regions in the amygdala and the thalamus. Injury to these critical structures also damages the brain circuitry and compromises the functionality housed in these brain regions. In agreement with these findings, widespread neuronal damage to structures in the limbic system and cortical areas was noted following DFP-SE (Figure 5). Sex-specific and regional differences in vulnerability to neuronal damage following DFP-SE have been previously reported. For example, one study reported that FJB+ cells were significantly higher in the dentate gyrus but not in the cortical region in adult female rats compared to adult male rats^74^. Interestingly, male juvenile rats exhibited greater cell injury across all brain regions following DFP-SE compared to female juvenile rats ^75^.

In our study, FJC+ cells were significantly higher in male rats in the CA1, DG, and PC regions. In contrast, female rats exhibited greater neuronal injury in the CA3 region following DFP-SE. VPM intervention had a significant neuroprotective action in the CA1 and PC regions for both sexes. However, VPM was protective in the DG region in only male rats. In the CA3 regions, VPM provided some neuroprotection, as evidenced by a lower number of FJC+ cells in both sexes following DFP-SE; however, significance could not be achieved. It is worth noting that male rats exhibited significantly higher VPM brain levels following DFP-SE, but the drug persisted longer in females. Whether these PK-related differences contribute to the sex-related differences in neuronal injury outcomes following DFP-SE will be addressed through investigations of VPM dose and duration in a future study.

We also investigated the pathological effects of repeated i.m. administrations of VPM by conducting muscle histology at the injection site ^67, 68^. Our studies demonstrated that the i.m. treatment regimen with VPM did not produce any overt inflammation or other pathology at the injection sites at both acute and chronic time points after the end of the injection period. No sex differences were noted in muscle pathology following i.m. VPM administrations (Figure 4). While a few clusters of scattered infiltrates indicative of mild inflammation were noted at acute time points, the pathology scores between DFP rats treated with SAL or VPM were not different. At 3 months after the last VPM injection, the muscle pathology revealed some minimal infiltrates in some rats, while the majority had pathology scores of 0, indicating the absence of inflammation. These studies establish the safety of i.m. VPM and demonstrates that this route of administration may be an effective method to deliver VPM post-SE. Our VPM formulation had an extended shelf life and showed little degradation even after 200 days following constitution (Table 6). The stability and ease of VPM constitution for i.m. administration will afford benefits for stockpiling VPM formulations in the event it is further optimized as an OP countermeasure.

**Table 6.** Stability of VPM formulation.

| **Stability Time Point** | **% Labeled Claim** |
| --- | --- |
| Day 0 | 98.21 ± 0.88 |
| Day 90 | 108.54 ± 1.79 |
| Day 203 | 108.65 ± 0.55 |

The stability of the VPM formulation was evaluated under ICH QA1 (R2) guidelines [Stability Testing of New Drug Substances and Products] and in accordance with the USP VPM injection monograph (USP29-NF24). VPM formulation (10 mg/mL in sterile saline) was stored in glass bottles at 25°C ± 2°C and 60 ± 5% Relative Humidity. At the respective time points, three bottles were randomly chosen, and VPM levels were assessed via HPLC methods.

Our study had some limitations. First, we did not assess electrographic SE and instead assessed SE severity on the Racine scale. We have previously shown that rats displaying a Racine score of 4 and above reliably exhibit electrographic EEG patterns akin to SE^5, 50^. Additionally, other studies have shown that rats from mixed-sex cohorts housed in the same room yielded reproducible SE severity in both sexes and both EEG telemetry (surgery) and non-telemetry (non-surgery) groups ^76^. Second, we did not quantify neuroinflammatory changes in this study. The goal of this study was to validate the safety and neuroprotective efficacy of a VPM formulation that could be administered i.m. after a reasonable delay following DFP-SE. Studies are underway in our laboratory to investigate the anti-inflammatory effects of VPM and its underlying molecular mechanism. In conclusion, the data from this study demonstrate that adjunct treatment with VPM could provide effective countermeasures for extending neuroprotective efficacy and improving long-term neurological outcomes after OP intoxication.

## Acknowledgement

This work was supported by the Countermeasures Against Chemical Threats (CounterACT) Program, the National Institutes of Health (NIH) Office of the Director (OD), and the National Institute of Neurological Disorders and Stroke (NINDS) [Grant Number-UG3 NS133630]. Its contents are solely the responsibility of the authors and do not necessarily represent the official views of the federal government. Services in support of the research project were generated by the VCU Massey Comprehensive Cancer Center Tissue and Data Acquisition and Analysis Shared Resource, supported, in part, with funding from NIH-NCI Cancer Center Support Grant P30 CA016059.

## Notes

### Competing Interest Statement

The authors have declared no competing interest.

